# Mutations in *ampD* cause hyperproduction of AmpC and CphA beta-lactamases and high resistance to beta-lactam antibiotics in *Chromobacterium violaceum*

**DOI:** 10.1101/2024.12.20.629803

**Authors:** Luís G. Laranjeiro, Carlos E. M. Neme, Maristela Previato-Mello, Bianca B. Batista, Isabel Henriques, José F. da Silva Neto

## Abstract

Bacterial resistance to beta-lactam antibiotics mediated by beta-lactamase enzymes is widespread worldwide. *Chromobacterium violaceum*, an environmental Gram-negative bacteria pathogen, is intrinsically resistant to some beta-lactam antibiotics. In this work, we found that mutations in an *ampD* gene, encoding a peptidoglycan-recycling amidase, cause hyperproduction of two chromosomal beta-lactamases (AmpC and CphA), conferring high beta-lactam resistance in *C. violaceum*. Susceptibility tests using Δ*ampC*, Δ*cphA*, and *ΔampC*Δ*cphA* mutant strains revealed specific susceptibility profiles to penicillin, cephalosporin, and carbapenem beta-lactams, suggesting that AmpC is a broad-spectrum beta-lactamase (penicillinase and cephalosporinase), while CphA is a narrow-spectrum metallo-carbapenemase. Beta-galactosidase assays indicate that the expression of *ampC* and *cphA* increased in response to beta-lactams. We isolated *C. violaceum* spontaneous mutants resistant to the antibiotic ceftazidime and found that most mutants were also resistant to several other beta-lactams and overexpressed *ampC* and *cphA*. DNA sequencing of the three paralog genes encoding the *C. violaceum* AmpD amidases revealed mutations of different types in AmpD1 (CV_0566) in most of the spontaneous mutants, but no mutation was found in AmpD2 or AmpD3. Analysis of single and combined null amidase mutants revealed overexpression of both beta-lactamases and increased resistance to beta-lactams only in mutants with deleted *ampD1*. When introduced into *ampD1* null or spontaneous mutants, the *ampD1* gene rescued the antibiotic-related phenotypes. The AmpD1 amidase from *C. violaceum* has a unique architecture with an N-terminal acetyltransferase domain. Our work offers new insights into the mechanisms of beta-lactamase-mediated antibiotic resistance and open perspectives to improve the treatment of *C. violaceum* infections.

## INTRODUCTION

Antimicrobial resistance has emerged and spread among bacterial pathogens, posing a threat to public health worldwide (1, 2). Resistance to beta-lactams, an important class of bactericidal antibiotics that block cell wall synthesis, is typically mediated by the beta-lactamases. These enzymes hydrolyze beta-lactams such as penicillin, cephalosporin, monobactam, and carbapenem with variable efficiency, using a conserved serine residue (classes A, C, and D beta-lactamases) or zinc as a cofactor (class B metallo-beta-lactamases) (3–5). Chromosomally encoded beta-lactamases such as AmpC (a class C cephalosporinase) (6) and CphA (a class B narrow-spectrum carbapenemase) (7–9) are important determinants of beta-lactam resistance in many bacterial pathogens, including *Pseudomonas aeruginosa*, *Stenotrophomonas maltophilia*, *Aeromonas spp*., and several bacteria from the *Enterobacteriaceae* family (6, 10).

Beta-lactam resistance mediated by chromosomal beta-lactamases is closely associated with perturbations in the pathways of peptidoglycan (PGN) synthesis and recycling that culminate in a high expression of beta-lactamase genes (10). In the case of transient induction, the presence of beta-lactams increases the products of the PGN metabolism, which are sensed by regulatory systems and transcription factors, such as AmpR and BlrAB, culminating in the activation of *ampC*, *cphA*, and other beta-lactamase genes (6, 10). Stable overexpression of these beta-lactamase genes is frequently associated with mutations in genes of the PGN recycling pathway. For instance, mutations in the *ampD* gene, which encodes the PGN amidase AmpD, are commonly found in several bacterial clinical isolates that are resistant to beta-lactam antibiotics by hyperproducing AmpC and other chromosomal beta-lactamase enzymes (6, 10–13).

*Chromobacterium violaceum*, a Gram-negative beta-proteobacterium commonly found in soil and water worldwide, is an environmental pathogen that causes serious infections in humans and other animals (14). Although rare, *C. violaceum* infections in humans show a rapid clinical course and have a high mortality rate, with bacteria spreading to several organs, including liver and spleen (14–17). Misdiagnosis and incorrect antibiotic prescription contribute to the unfavorable outcome, given that many clinical *C. violaceum* isolates are intrinsically resistant to some antibiotics, including beta-lactams (18–21).

Chromosomally encoded beta-lactamases have been studied in many environmental opportunistic pathogens (6, 10), but few studies have investigated the molecular mechanisms of antibiotic resistance in *Chromobacterium* species (22, 23). Genome sequence analysis has revealed the presence of two predicted chromosomal beta-lactamase genes, *ampC* and *cphA*, in *C. violaceum* ATCC 12472 (24, 25) and in clinical *C. violaceum* isolates (26, 27), but their contribution to beta-lactam resistance in *C. violaceum* remains to be determined. In this work, we investigated the role and regulation of *ampC* and *cphA* in response to beta-lactams. We found that these genes confer resistance to distinct beta-lactams and their hyperproduction arise from mutations in the amidase AmpD1.

## RESULTS

### Two chromosomal beta-lactamases of distinct classes confer high resistance to beta-lactam antibiotics in *C. violaceum*

Many clinical *C. violaceum* strains are resistant to beta-lactam antibiotics (19, 20), but the mechanisms of resistance are unknown. The genome of *C. violaceum* ATCC 12472 has two genes, CV_1310 and CV_3150, which were predicted to encode chromosomal beta-lactamase enzymes (24, 25). Our *in silico* analysis with the amino acid sequence alignment using BlastP revealed that outside the *Chromobacterium* genus, the best hits were AmpC from *Pseudomonas aeruginosa*, showing 60% similarity with CV_1310, and CphA from *Aeromonas hydrophila*, showing 65% similarity with CV_3150. Therefore, we opted to refer to these beta-lactamases as AmpC and CphA in *C. violaceum*.

We constructed Δ*ampC*, Δc*phA*, and Δ*cphA*Δ*ampC* null mutant strains and compared their beta-lactam resistance profile with that from the wild type (WT). Resistance was evaluated through minimum inhibitory concentration (MIC) assays (Table 1) and disk diffusion tests (Figure 1A and B), and the mutant phenotypes were further validated by genetic complementation. A Δ*ampC* mutant strain showed increased sensitivity to penicillin, cephalosporin, and monobactam beta-lactams but not to carbapenems. On the other hand, a Δ*cphA* mutant strain was sensitive to the two tested carbapenems but not to the other beta-lactams. A Δ*cphA*Δ*ampC* double mutant strain showed the phenotypes observed in each individual mutant strain (Figure 1A and Table 1). Complementation by providing the *ampC* or *cphA* genes into each mutant restored or even increased the resistance to the beta-lactams, while an empty vector had no effect (Figure 1B and Table 1). When the plasmid carrying either the *ampC* or the *cphA* gene was introduced into *Escherichia coli* DH5α, the level of resistance to the same beta-lactams was barely increased (Figure S1). To investigate whether AmpC or CphA are metallo-carbapenemases, we performed mCIM and eCIM tests using imipenem (28). WT *C. violaceum* showed a metal-dependent carbapenemase activity, which was abolished in a Δ*cphA* but unaffected in a Δ*ampC* strain. As a control, a *Klebsiella pneumoniae* reference strain presented a metal-independent carbapenemase activity (Figure 1C). Altogether, these data indicate that CphA is a metallo-beta-lactamase conferring resistance to carbapenems, while AmpC is a broad-spectrum beta-lactamase that confers resistance to penicillin and cephalosporin beta-lactams.

**Figure 1.**
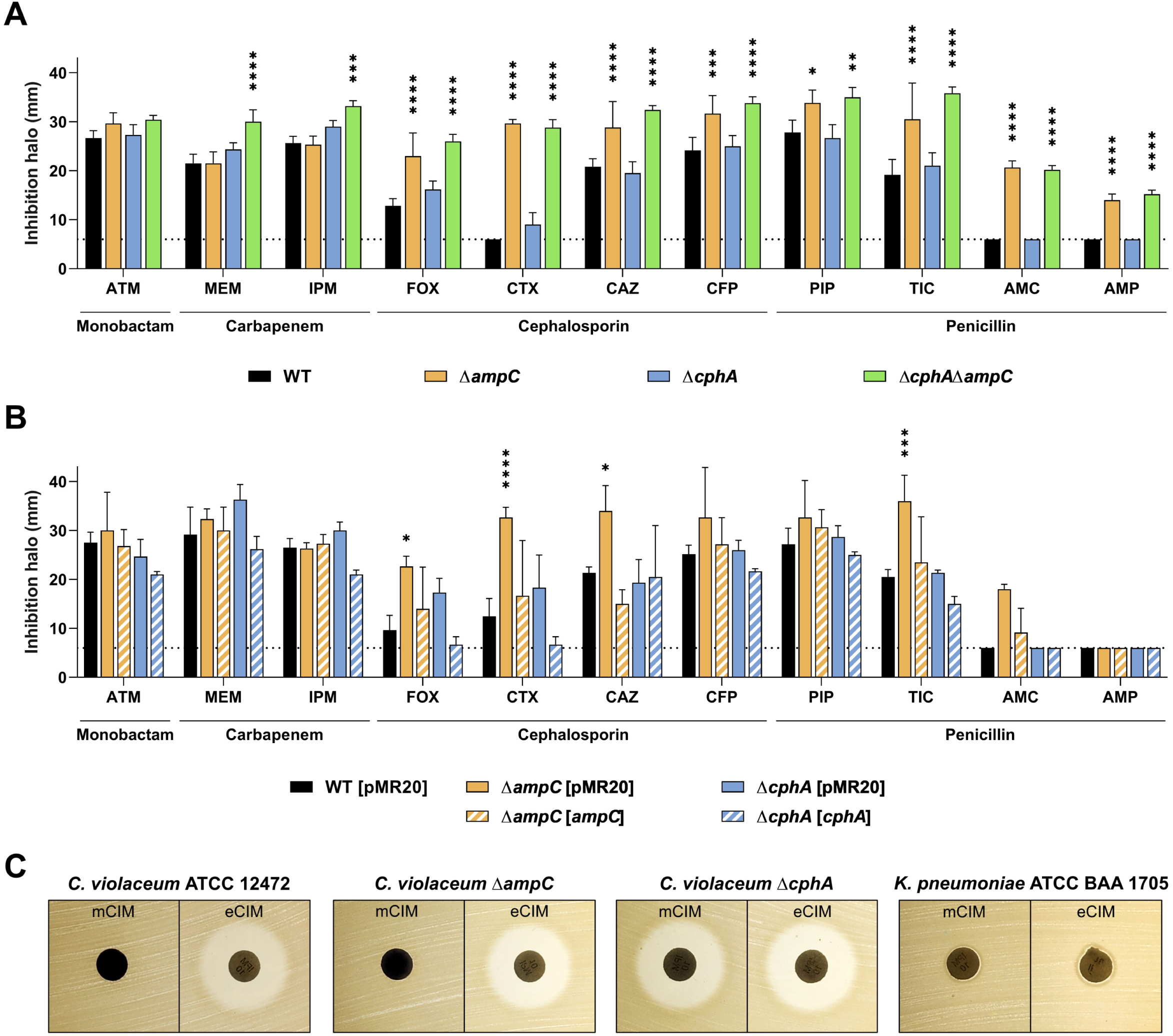
*C. violaceum* harbors two active beta-lactamases. (A and. **B)** Disk-diffusion assays. **(A)** Deletion of *ampC* and *cphA* genes increases *C. violaceum* susceptibility to different beta-lactam antibiotics. **(B)** Complementation restores the phenotype of the mutants to a level similar to the wild type strain. *C. violaceum* WT with and without the empty pMR20 vector were used as controls. Inhibition halos are shown in mm. Dotted lines indicate the diameter of the disks (6 mm). ATM, aztreonam; MEM, meropenem; IPM, imipenem; FOX, cefoxitin; CTX, cefotaxime; CAZ, ceftazidime; CFP, cefoperazone; PIP, piperacillin; TIC, ticarcillin; AMC, amoxicillin-clavulanic acid; and AMP, ampicillin. Charts representing the average of the halos measured between the experimental triplicate. ****p < 0.0001; ***p < 0.001; **p < 0.01; * p < 0.05. Two-way ANOVA followed by Tukey’s multiple comparisons test. **(C)** mCIM (modified Carbapenem Inactivation Method) and eCIM (EDTA-modified Carbapenem Inactivation Method) tests indicate that CphA is an active metallo-beta-lactamase (MBL).

**Table 1.** Antibiotic resistance profile of *C. violaceum* beta-lactamase mutants.

| STRAIN | MIC (µg/mL) |  |  |  |  |  |  |  |
| --- | --- | --- | --- | --- | --- | --- | --- | --- |
|  | MEM | IPM | CAZ | CTX | FOX | AMP | AMC | TZP |
| WT | 0.25 | 2 | 32 | 512 | 64 | 1024 | 1024 | 16 |
| WT [pMR20] | 0.25 | 2 | 32 | 512 | 64 | 1024 | 1024 | 16 |
| $\Delta ampC$ | 0.25 | 2 | 1 | 2 | 8 | 16 | 16 | 2 |
| $\Delta ampC$ [pMR20] | ND | ND | 2 | 4 | ND | 32 | 32 | 4 |
| $\Delta ampC$ [*ampC*] | 0.25 | 2 | 128 | 512 | 256 | 1024 | 1024 | 16 |
| $\Delta cphA$ | 0.125 | 1 | 32 | 256 | 32 | 512 | 512 | 8 |
| $\Delta cphA$ [pMR20] | 0.125 | 1 | 64 | 256 | 64 | 512 | 512 | 8 |
| $\Delta cphA$ [*cphA*] | 4 | 8 | 128 | 512 | 256 | 1024 | 1024 | 16 |
| $\Delta cphA\Delta ampC$ | 0.125 | 1 | 1 | 2 | 8 | 16 | 16 | 1 |

ND = Not determined; WT, wild type; SM, spontaneous mutant; MEM, meropenem; IPM, imipenem; CAZ, ceftazidime; CTX, cefotaxime; FOX, ceftoxitin; AMP, ampicillin; AMC, amoxicillin-clavulanic acid; TZP, piperacillin-tazobactam.

### Both *C. violaceum* beta-lactamases are induced by beta-lactam antibiotics

Some chromosomal beta-lactamases are induced in the presence of different beta-lactam antibiotics (6, 29). To check whether this is the case for *C. violaceum* beta-lactamases, we cloned the promoter regions of *ampC* (P*ampC*) and *cphA* (P*cphA*) into a *lacZ*-based reporter vector. The resulting constructs were introduced into the *C. violaceum* WT strain. The promoter activities were quantified by beta-galactosidase assays from midlog phase-cultures treated with different antibiotics (Figure 2). The expression of *ampC* and *cphA* increased in the presence of beta-lactam antibiotics, but while ceftazidime (CAZ) acted as a weak inducer, imipenem (IPM), ampicillin (AMP), and cefoxitin (FOX) were strong inducers (Figure 2). In cultures treated with eight non-beta-lactam antibiotics of distinct classes, the promoters of *ampC* and *cphA* showed a basal expression, comparable with that of untreated cultures (Figure 2). These results demonstrate that in *C. violaceum*, the expression of both beta-lactamases increases when exposed to beta-lactam antibiotics.

**Figure 2.**
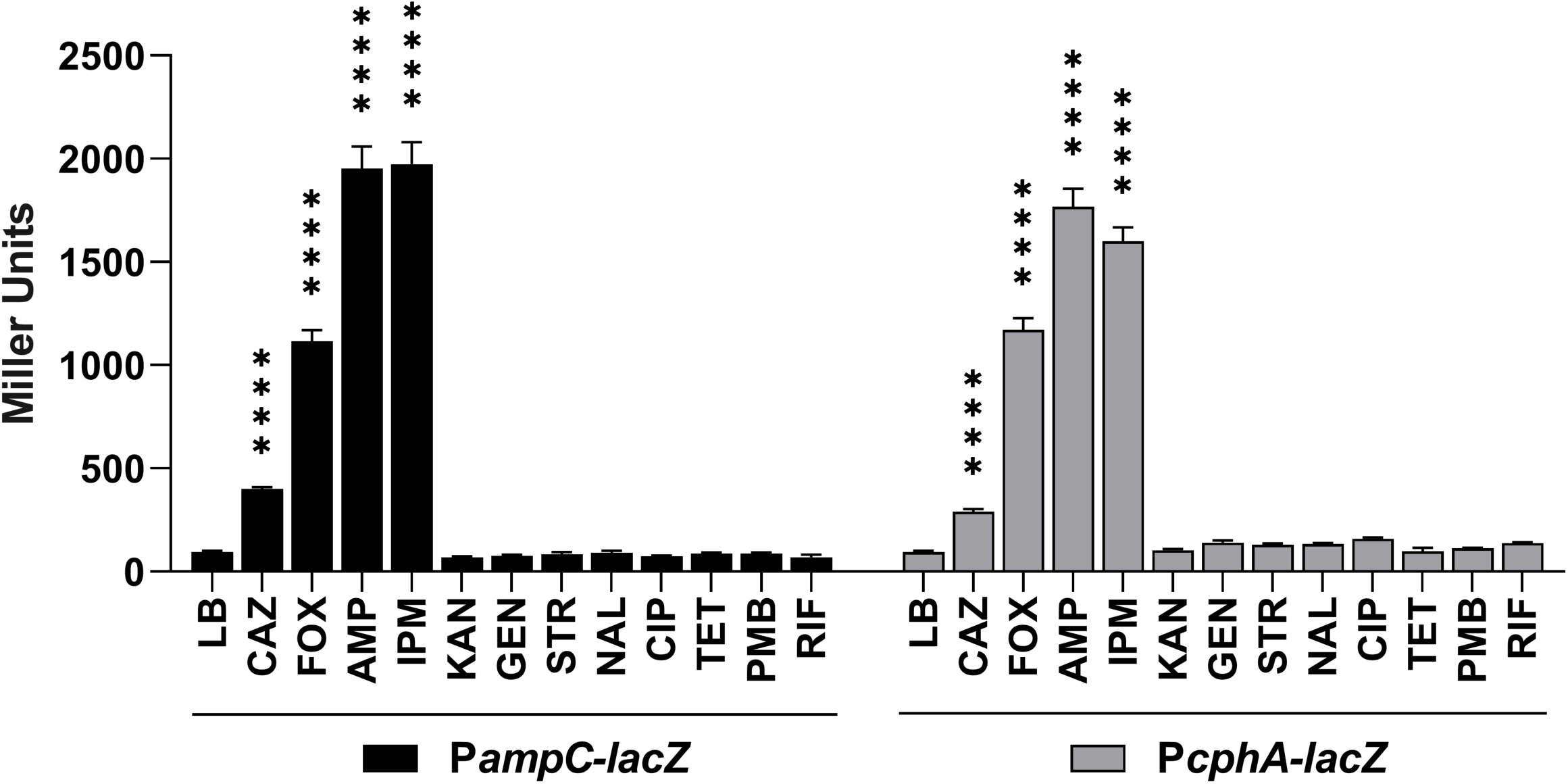
Beta-lactam antibiotics induce *ampC* and *cphA* beta-lactamases. The expression was measured by beta-galactosidase activity assay. *C. violaceum* WT harboring the indicated *lacZ* fusions was cultured in LB or LB plus the following antibiotics: CAZ, ceftazidime; FOX, cefoxitin; AMP, ampicillin; IPM, imipenem; KAN, kanamycin; GEN, gentamicin; STR, streptomycin; NAL, nalidixic acid; CIP, ciprofloxacin; TET, tetracycline; PMB, polymyxin B; and RIF, rifampicin. The error bars represent the standard deviation of the mean of biological quintuplicate. Stars indicate statistical significance compared to the wild type strain. ****p < 0.0001; ***p < 0.001; **p < 0.01; * p < 0.05. Two-way ANOVA followed by Tukey’s multiple comparisons test.

### Spontaneous mutants isolated in ceftazidime are resistant to various beta-lactams without loss of fitness

To identify mutations associated with beta-lactam resistance in *C. violaceum*, we isolated spontaneous mutants by plating overnight cultures of the WT strain ATCC 12472 on Mueller Hinton (MH) agar supplemented with increasing concentrations of CAZ. We chose this cephalosporin because it is a weak inducer of beta-lactamases in *C. violaceum* (Figure 2) and in other bacteria as well (12, 30, 31). A total of 60 colonies were isolated on MH plates containing 80 and 160 μg/mL of CAZ and named SM1 to SM60 (SM stands for spontaneous mutant). After replating the isolates on antibiotic-free MH and determine the MIC by the agar-dilution assay, 13 SM isolates showing MIC values greater than the control were selected (Table 2). We tested the beta-lactam resistance profile of the 13 selected SM isolates by disk diffusion using disks of nine beta-lactam antibiotics (Figure 3). The isolates SM1 and SM2 were highly resistant to all tested antibiotics. All the other SM isolates showed increased resistance to most of the tested beta-lactams, except for the carbapenems (Figure 3). These data indicate that the 13 spontaneous mutants isolated in CAZ are resistant to several beta-lactam antibiotics.

**Figure 3.**
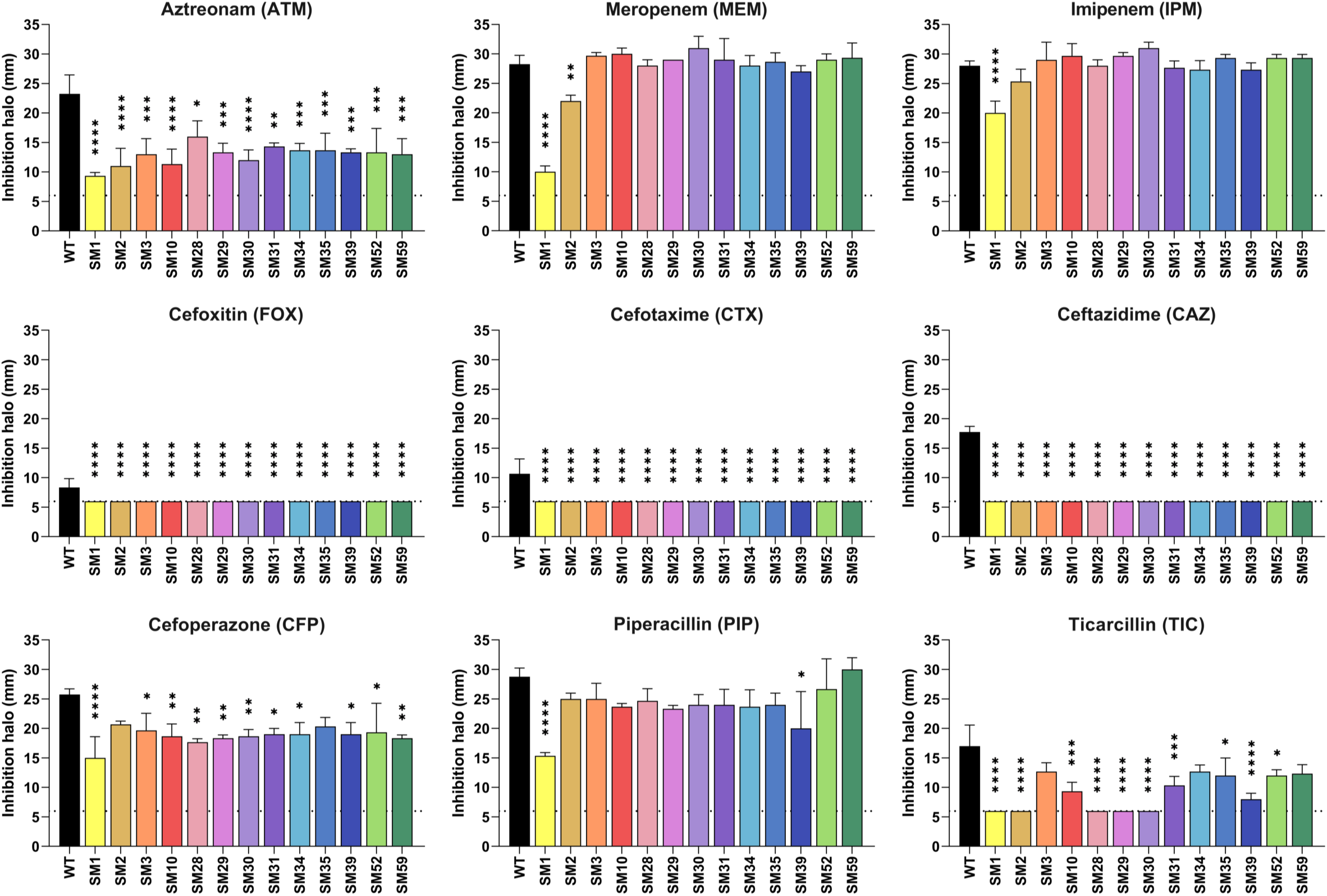
*C. violaceum* spontaneous mutants isolated in ceftazidime are resistant to various beta-lactams. Resistance profile of the SM isolates against different beta-lactam antibiotics by disk-diffusion assay. Average measurements of the halos from triplicate samples are shown. Halo inhibition shown in mm. Dotted lines indicate the diameter of the disks (6 mm). Stars indicate statistical significance compared to the wild type strain. ****p < 0.0001; ***p < 0.001; **p < 0.01; * p < 0.05. One-way ANOVA followed by Tukey’s multiple comparisons test.

**Table 2.** Ceftazidime MIC of *C. violaceum* wild type (WT) and spontaneous mutants (SM).

| STRAIN | MIC CAZ (µg/mL) |
| --- | --- |
| WT | 32 |
| SM1 | 1024 |
| SM2 | 512 |
| SM3 | 512 |
| SM10 | 512 |
| SM28 | 512 |
| SM29 | 512 |
| SM30 | 512 |
| SM31 | 512 |
| SM34 | 512 |
| SM35 | 512 |
| SM39 | 512 |
| SM52 | 512 |
| SM59 | 512 |

We assessed the fitness of the SM isolates by measuring their growth and survival in LB medium (Figure S2). In general, the SM isolates had a growth profile similar to that of the WT strain, except for SM1, which showed a slight delay, reaching the stationary phase later than the other strains (Figure S2A). Despite this, SM1 showed no change in viability after 20 hours of cultivation in LB medium, while the other isolates showed little or no change in viability (Figure S2B). Therefore, these data indicate that despite showing resistance to various beta-lactam antibiotics, we did not observe a significant disturbance in fitness in any of the 13 SM isolates.

### Except for SM1, all spontaneous mutants overexpress both beta-lactamases

To associate the beta-lactam antibiotic resistance profile of the SM isolates with the expression of the AmpC and CphA beta-lactamases, we carried out beta-galactosidase activity assays using the P*ampC* and P*cphA* transcriptional fusions in the SM isolates (Figure 4). Except for SM1, the *ampC* and *cphA* promoters were highly expressed in all SM isolates in comparison with the basal activity found in the WT. The SM2 isolate showed a modest expression compared with the high expression of the other SM isolates (Figure 4A and B). To check that the *lacZ* fusions were functional in the SM1 isolate, the antibiotic AMP was added to the cultures for 30 minutes. In such condition, both promoters were highly induced in both the WT and SM1 strains (Figure 4C). These data indicate that in 12 of the 13 SM isolates, the increased resistance to beta-lactam antibiotics can be attributed to the simultaneous overexpression of the *ampC* and *cphA* beta-lactamases. The high resistance of SM1 to beta-lactams does not seem to be linked to an altered expression of the AmpC or CphA beta-lactamases.

**Figure 4.**
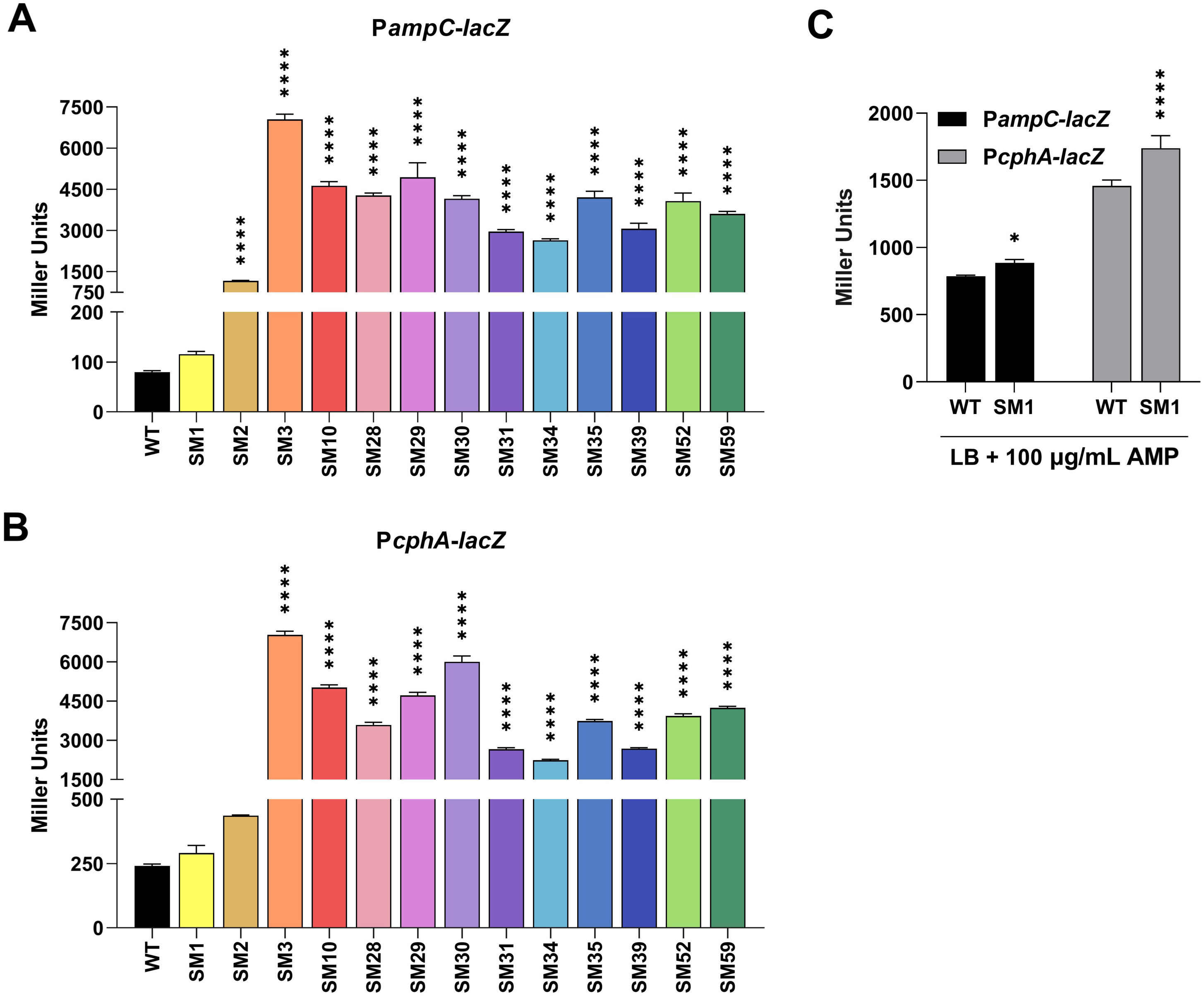
Almost all spontaneous mutants overexpress both beta-lactamases. (A and. **B)** Promoter activity of both beta-lactamases in LB, and **(C)** LB in the presence of 100 μg/mL AMP. Beta-galactosidase assays were carried out on WT and SM isolates harboring the P*ampC-lacZ* or P*cphA-lacZ* fusions. The error bars represent the standard deviation of the mean of a biological quintuplicate. Stars indicate statistical significance compared to the wild type strain. ****p < 0.0001; ***p < 0.001; **p < 0.01; * p < 0.05. One-way ANOVA followed by Tukey’s multiple comparisons test.

### Except for SM1 and SM2, all spontaneous mutants have mutation in the *ampD1* **gene**

Given the importance of the amidase AmpD in the peptidoglycan recycling process and its close relationship with beta-lactam resistance (12, 32, 33), we investigated the occurrence of mutations in *ampD* in the SM isolates. We found three AmpD paralogs in *C. violaceum* that we named as *ampD1*, *ampD2*, and *ampD3*. In the genome, the gene *ampD2* is located next to *ampC* (Figure 5A). The three AmpD proteins shared 30 to 40% similarity (Figure 5B) and have an amidase-2 domain (AMI_2; PF01510) (Figure 5C), found in *E. coli* (AmpD and AmiD) and *P. aeruginosa* (AmpD, AmpDh2, and AmpDh3) (34). Interestingly, *C. violaceum* AmpD1 has an acetyltransferase domain (ACTF_1) at the N-terminus (Figure 5C), which was not annotated in the original version of the genome. A potential fourth amidase in *C. violaceum*, CV_3822 (AmpD4), possesses an amidase-3 domain (AMI_3; PF01520) and shows low similarity to the other three AmpDs (18 to 25% in full-length and 15 to 19% in amidase domain alignments) (Figure 5C).

**Figure 5.**
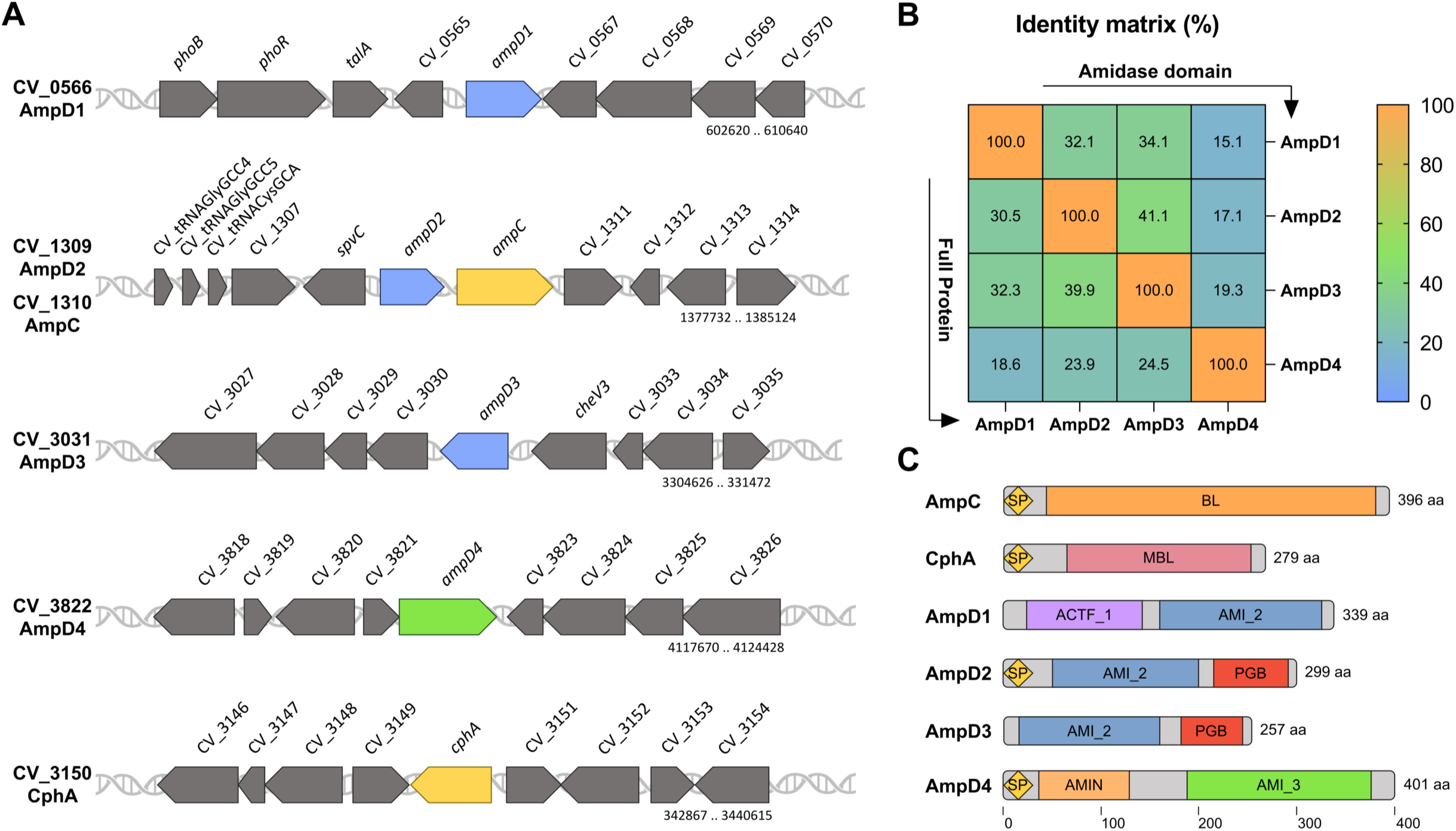
Genomic organization, domain architecture, and multiple alignment of *C. violaceum* amidases. **(A)** Genomic map of the genes encoding beta-lactamases and AmpD amidases. **(B)** Similarity matrix between the four *C. violaceum* amidases based on multiple amino acid alignment using Clustal Omega. Alignments were performed using the full-length proteins or only their amidase domians. **(C)** Domain architecture of the amidases. Each color indicates a domain: SP, signal peptide; BL, beta-lactamase; MBL, metallo-beta-lactamase; ACTF_1, type 1 acetyltransferase; AMI_2, type 2 amidase; PGB, peptidoglycan-binding; AMI_3, type 3 amidase; AMIN, Amidase N-terminal domain. The genes have the following entries in KEGG, NCBI, and UniProt databases: AmpD1, CV_0566, CV_RS02775, Q7P0K1; AmpD2, CV_1309, CV_RS06380, Q7NYG5; AmpD3, CV_3031, CV_RS23055, Q7NTM3; AmpD4, CV_3822, CV_RS18925, Q7NRF9; AmpC, CV_1310, CV_RS06385, Q7NYG4; CphA, CV_3150, CV_RS15465, Q7NTA9.

The paralog genes *ampD1*, *ampD2*, and *ampD3* were sequenced by the Sanger method using DNA from the *C. violaceum* ATCC 12472 as a control and from the 13 SMs isolated in CAZ. The DNA sequences were compared by BLAST against the reference genome of the *C. violaceum* ATCC 12472 strain (24). The alignment shows no mutations in the three *ampD* genes in the WT strain, as expected. No mutations were found in the *ampD2* and *ampD3* genes in any of the SM isolates. Except for SM1 and SM2, several mutations were detected in the *ampD1* gene in all SM isolates (Table 3; Figure 6). Nonsense mutations were detected in the SM3, SM10, SM52, and SM59 isolates that replaced tryptophan codons with a stop codon (Table 3). These mutations occurred in the N-terminal acetyltransferase domain where they generate truncated versions of inactive AmpD1 proteins (Figure 6). Another six SM isolates showed missense mutations inside the amidase domain, while the SM30 had a *frameshift* due to an 11 bp deletion that generates a stop codon in the middle of the *ampD1* gene (Table 3; Figure 6). Comparing the mutation profile observed in *ampD* mutants from other bacteria with that found in *ampD1* from *C. violaceum*, we discovered many mutations that have already been described and 5 novel mutations at different regions of the protein (Figure 6A). Collectively, our data indicate that out of the 13 SMs isolated in CAZ, 12 overexpressed *ampC* and *cphA* and 11 had point mutations in the *ampD1* gene (CV_0566). These data suggest that AmpD1 plays a greater role than the other two AmpD paralogs in beta-lactam resistance in *C. violaceum*.

**Figure 6.**
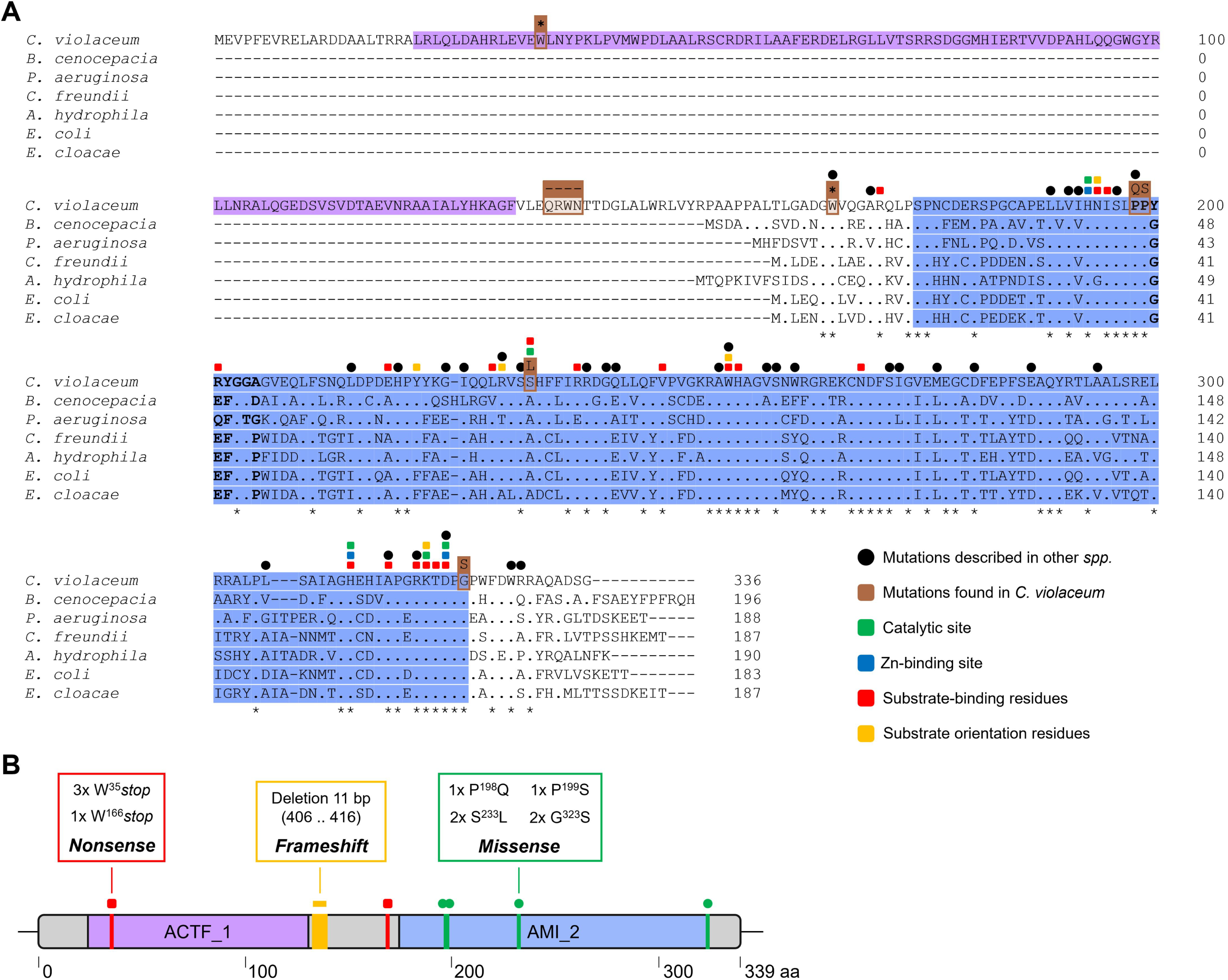
Sequence alignment of AmpD amidases. **(A)** Multiple amino acid alignment performed with Clustal Omega. AmpD1 from *Chromobacterium violaceum* ATCC 12472 (Q7P0K1_CHRVO) was aligned with AmpD sequences from different bacteria, showing the following identity percentages: *Burkholderia cenocepacia* J2315 (B4E5W0_BURCJ, 57%), *Pseudomonas aeruginosa* PAO1 (G3XCW9_PSEAE, 55%), *Citrobacter freundii* OS60 (AMPD_CITFR, 53%), *Aeromonas hydrophila* ATCC 7966 (A0KPV3_AERHH, 52%), *Escherichia coli* K12 (AMPD_ECOLI, 52%), and *Enterobacter cloacae* strain 14 (AMPD_ENTCL, 51%). The protein domains are marked in purple (acetyltransferase) and blue (type 2 amidase). Conserved residues are indicated by colored squares. Black circles indicate mutations reported in other bacteria. The brown boxes indicate mutations found in *C. violaceum ampD1*: replaced amino acids shown at the top; dashes indicate a *frameshift* event, and the star indicates the stop codon. Residues in bold correspond to the loop of the conserved r2 region in the amidase domain responsible for changing the inactive and active state. **(B)** Summary of mutations found in *ampD1* in *C. violaceum* spontaneous CAZ-resistant mutants.

**Table 3.** Mutations in *ampD1* of CAZ spontaneous mutants.

| STRAIN | MUTATION | ALTERATION | POSITION | CLASSIFICATION |
| --- | --- | --- | --- | --- |
| **Deletion – *ampD1* (CV_0566)** |  |  |  |  |
| SM30 | Frameshift | -11 pb | (406 .. 416) |  |
| **Substitutions – *ampD1* (CV_0566)** |  |  |  |  |
| SM10 | Nonsense | TGG → TAG | W35* | Aromatic → Stop codon |
| SM52 | Nonsense | TGG → TAG | W35* | Aromatic → Stop codon |
| SM59 | Nonsense | TGG → TAG | W35* | Aromatic → Stop codon |
| SM3 | Nonsense | TGG → TAG | W166* | Aromatic → Stop codon |
| SM28 | Missense | CCG → CAG | P198Q | NP/Aliphatic → PNC |
| SM34 | Missense | CCT → TCT | P199S | NP/Aliphatic → PNC |
| SM29 | Missense | TCG → TTG | S233L | PNC → NP /Aliphatic |
| SM35 | Missense | TCG → TTG | S233L | PNC → NP/Aliphatic |
| SM31 | Missense | GGT → AGT | G323S | NP/Aliphatic → PNC |
| SM39 | Missense | GGT → AGT | G323S | NP/Aliphatic → PNC |

Analysis carried out on the NCBI database with the Blast tool, using the Blastn and Blastx options. In bold are the bases changed by the mutations. SM: spontaneous mutant; \*: stop codon; PNC: polar non-charged; NP: nonpolar.

### Mutations in *ampD1*, but not in *ampD2* and *ampD3*, increase resistance to beta-lactams via overexpression of beta-lactamases in *C. violaceum*

To further investigate the role of the three *C. violaceum* AmpD paralogs in beta-lactam resistance, we constructed single null mutants Δ*ampD1*, Δ*ampD2*, and Δ*ampD3*, double mutants Δ*ampD1D2*, Δ*ampD1D3*, and Δ*ampD2D3*, and a triple mutant Δ*ampD1D2D3*. Disk diffusion assays indicated that only mutants with *ampD1* deleted (Δ*ampD1*, Δ*ampD1D2*, Δ*ampD1D3*, and Δ*ampD1D2D3*) had increased resistance to the tested beta-lactam antibiotics, if compared with the WT. None of the mutant strains showed a resistance phenotype to the tested carbapenems (IPM and MEM) (Figure 7). These data are consistent with the beta-lactam resistance phenotypes observed for the SM isolates harboring point mutations in *ampD1* (Figure 3). We also inserted the *ampD1* gene cloned into a vector in Δ*ampD1* and all 11 SM isolates harboring a point mutation in *ampD1*. In all these complemented strains, the increased resistance of the mutants to CAZ was rescued to patterns like those observed for the WT strain, except for SM59 (Figure 8). These data indicate that the increased resistance to beta-lactams in the *ampD1* null mutant and SM isolates (except SM59) is exclusively due to mutations in the *ampD1* gene.

**Figure 7.**
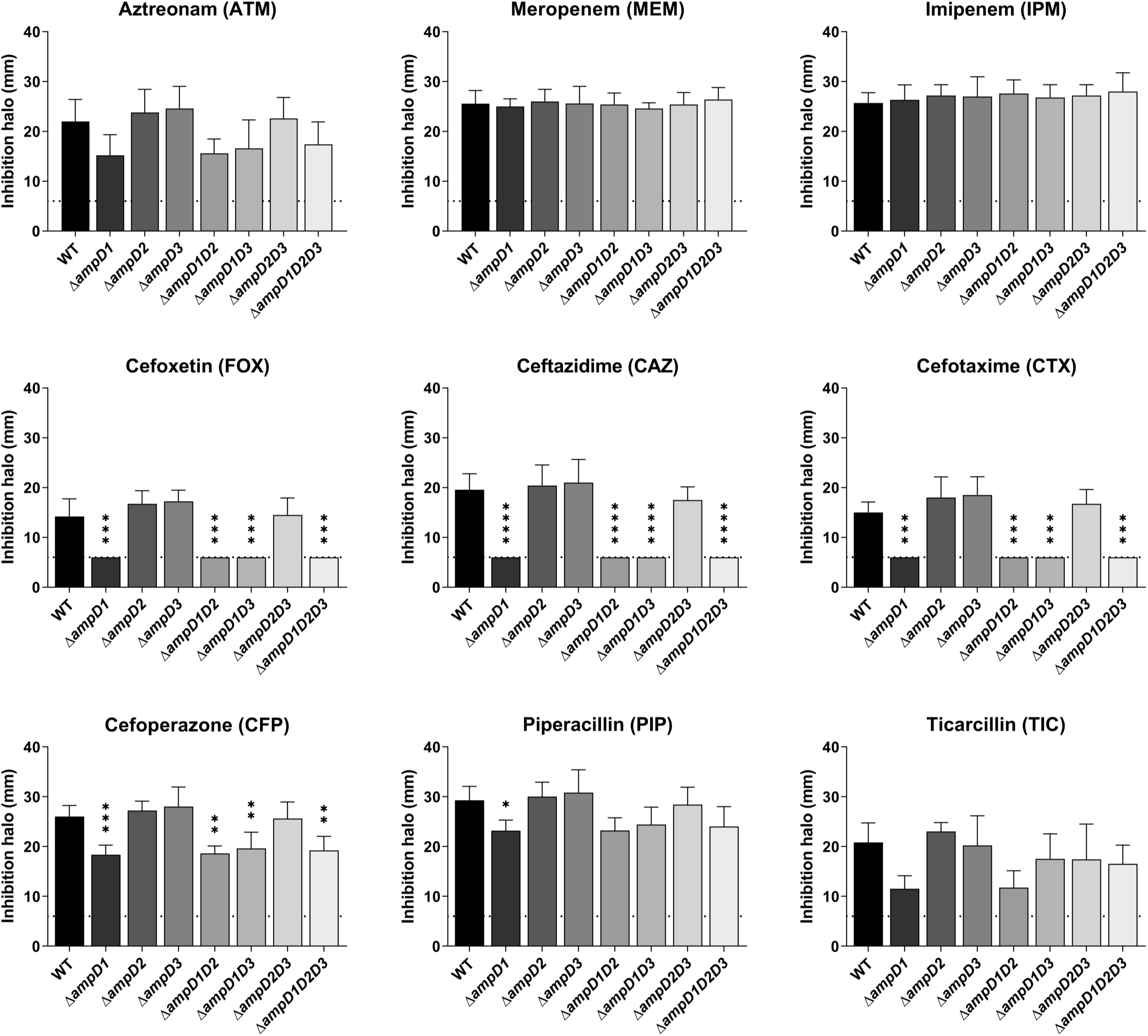
Null mutants with *ampD1* deletion show increased resistance to beta-lactams. Resistance profile of the amidase null mutants against different beta-lactam antibiotics by disk-diffusion assay. Average measurements of the halos from triplicate samples are shown. Halo inhibition shown in mm. Dotted lines indicate the diameter of the disks (6 mm). All the mutant strains were compared with the wild type strain. ****p < 0.0001; ***p < 0.001; **p < 0.01; * p < 0.05. One-way ANOVA followed by Tukey’s multiple comparisons test.

**Figure 8.**
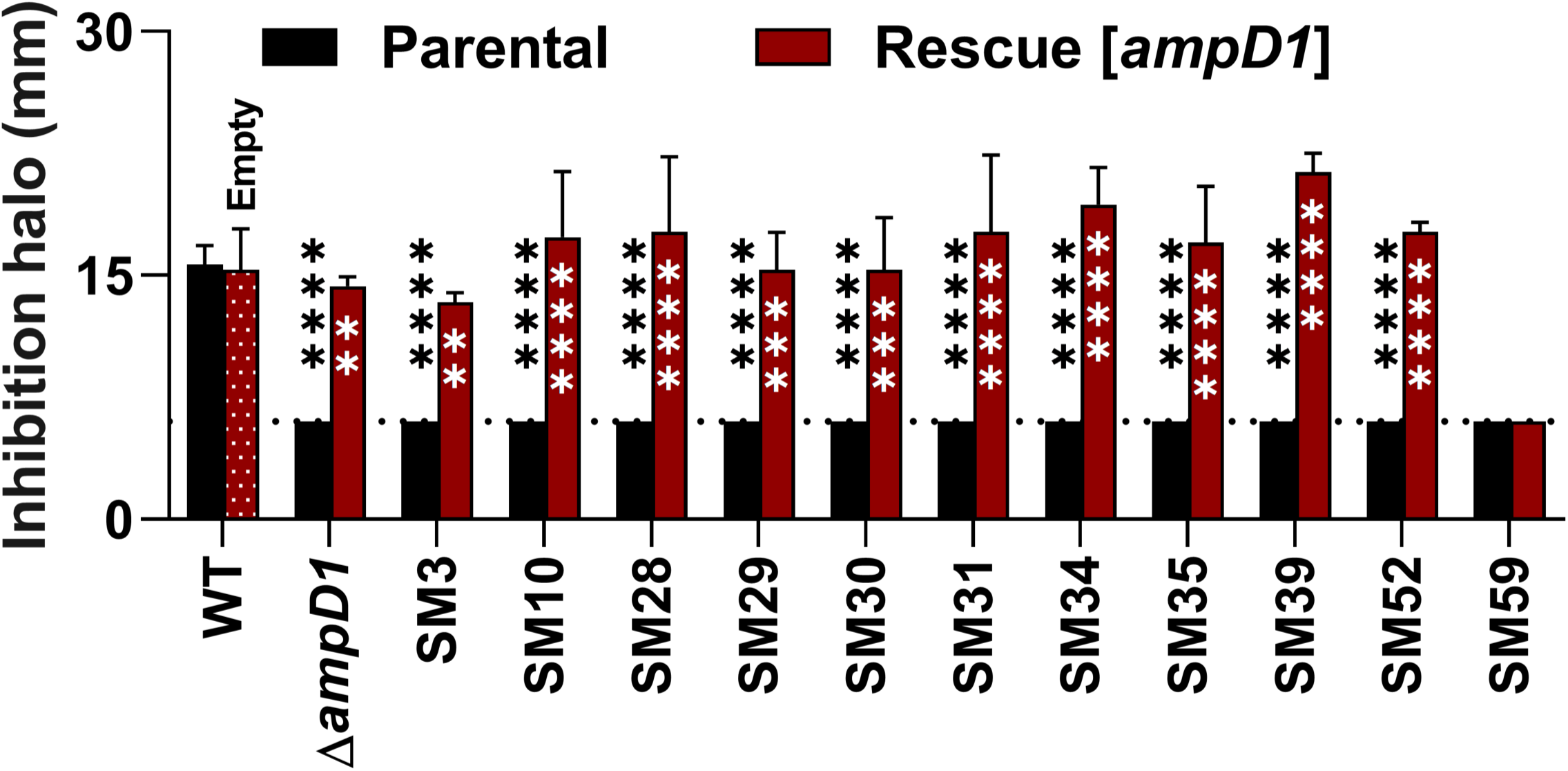
Complementation with the *ampD1* gene rescue the CAZ susceptibility of null and SM strains. The Δ*ampD1* null mutant and all SMs harboring mutation in *ampD1* (parental) were complemented with the *ampD1* gene cloned into the pMR20 vector (rescue). Disk-diffusion assays for CAZ were performed from triplicate samples. *C. violaceum* WT with and without the empty pMR20 vector were used as controls. Inhibition halos presented in mm. Dotted lines indicate the diameter of the disks (6 mm). Black asterisks indicate comparisons with the wild type strain, while white asterisks indicate the comparison in the same strain. ****p < 0.0001; ***p < 0.001; **p < 0.01; * p < 0.05. Two-way ANOVA followed by Tukey’s multiple comparisons test.

The agar-dilution test to determine the MIC was also carried out for some selected strains (Table 4). As expected from the previous tests, all the mutants with deleted *ampD1* showed resistance to most of the tested antibiotics, if compared with the WT strain, with MIC values at least 5 and 4 times higher for CAZ and FOX, respectively. No resistance phenotype was observed in the mutants of other amidases. A representative spontaneous mutant, the SM3 isolate (nonsense mutation W166stop in *ampD1*), showed MIC values similar to the Δ*ampD1* mutant. In all cases, by providing the *ampD1* gene but not the empty vector, the sensitivity phenotype was rescued (Table 4).

**Table 4.** Antibiotic resistance profile of *C. violaceum* null *ampD* mutants.

| STRAIN | MIC (µg/mL) |  |  |  |  |  |  |  |
| --- | --- | --- | --- | --- | --- | --- | --- | --- |
|  | MEM | IPM | CAZ | CTX | FOX | AMP | AMC | TZP |
| WT | 0.25 | 2 | 32 | 512 | 64 | 1024 | 1024 | 16 |
| WT [pMR20] | 0.25 | 2 | 32 | 512 | 64 | 1024 | 1024 | 16 |
| *ΔampD1* | 0.5 | 2 | 512 | 1024 | 512 | 1024 | 1024 | 16 |
| *ΔampD1* [pMR20] | 0.5 | 2 | 512 | 1024 | 512 | 1024 | 1024 | 16 |
| *ΔampD1* [*ampD1*] | 0.25 | 2 | 32 | 256 | 64 | 1024 | 1024 | 8 |
| *ΔampD2* | ND | ND | 32 | ND | ND | ND | ND | ND |
| *ΔampD3* | ND | ND | 32 | ND | ND | ND | ND | ND |
| *ΔampD1D2* | ND | ND | 512 | ND | ND | ND | ND | ND |
| *ΔampD1D3* | ND | ND | 512 | ND | ND | ND | ND | ND |
| *ΔampD2D3* | ND | ND | 32 | ND | ND | ND | ND | ND |
| *ΔampD1D2D3* | ND | ND | 512 | ND | ND | ND | ND | ND |
| SM3 | 0.5 | 2 | 512 | 1024 | 512 | 1024 | 1024 | 16 |
| SM3 [*ampD1*] | 0.25 | 1 | 128 | 256 | 32 | 512 | 128 | 8 |

ND = Not determined; WT, wild type; SM, spontaneous mutant; MEM, meropenem; IPM, imipenem; CAZ, ceftazidime; CTX, cefotaxime; FOX, cefoxitin; AMP, ampicillin; AMC, amoxicillin-clavulanic acid; TZP, piperacillin-tazobactam.

The expression of *ampC* and *cphA* was evaluated in the three *ampD-*null mutants, Δ*ampD1*, Δ*ampD2*, and Δ*ampD3* (Figure 9). Both P*ampC* and P*cphA* showed high beta-galactosidase activity in the Δ*ampD1* mutant, indicating overexpression of the two beta-lactamases (Figure 9A), which may explain the beta-lactam resistance phenotype in the null mutants and SMs harboring a mutation in the *ampD1* gene. Both promoters had only basal expression in Δ*ampD2* and Δ*ampD3*, comparable to that observed in the WT (Figure 9A), indicating that AmpD2 and AmpD3 are not involved with the expression of the beta-lactamases. The addition of AMP increased *ampC* and *cphA* expression in all cases, indicating that the reporter fusions in these strains were functional (Figure 9B). We also investigated the expression of *ampC* and *cphA* by RT-qPCR (Figure 9C and D). Consistently with beta-galactosidase assays, the SM2, SM59, and Δ*ampD1* mutants showed high expression of *ampC* and *cphA* in LB medium with or without AMP, indicating that these strains overexpress the beta-lactamases even in the absence of beta-lactams. In contrast, the SM1, Δ*ampD2*, and Δ*ampD3* strains showed an inducible expression pattern of *ampC* and *cphA*, as observed in the WT strain (Figure 9C and D).

**Figure 9.**
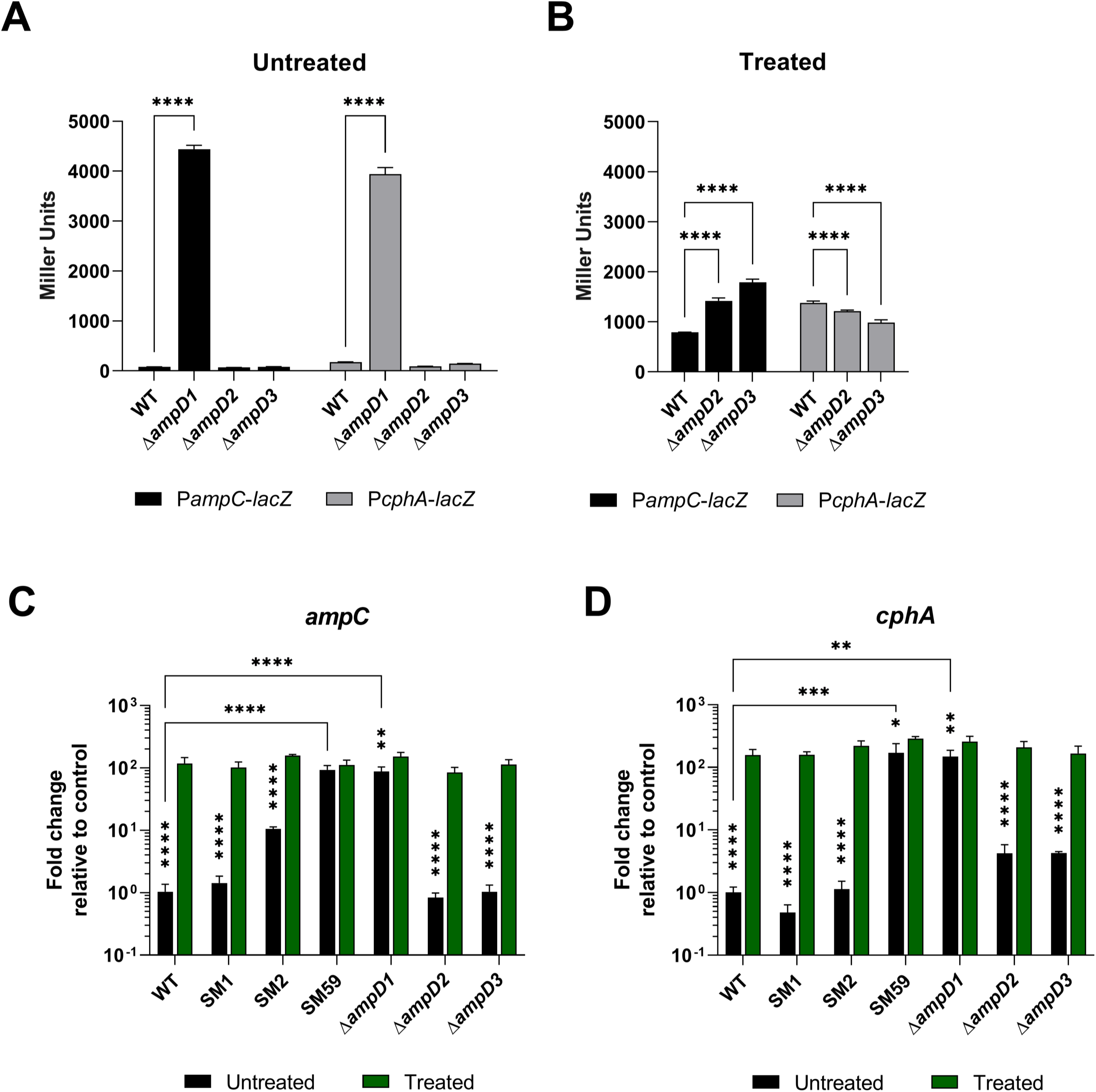
Deletion of *ampD1* increases the expression of beta-lactamases in *C. violaceum*. **(A)** The promoter activity of the beta-lactamase genes was evaluated in the indicated strains by beta-galactosidase activity assay. **(B)** Addition of 100 μg/mL AMP induces promoter expression beyond the basal level. **(C)** Analysis of *ampC* and **(D)** *cphA* mRNA expression in different strains in control condition (LB) and with 100 μg/mL AMP treatment by RT-qPCR. Assays were performed in biological triplicate. ****p < 0.0001; ***p < 0.001; **p < 0.01; * p < 0.05. Vertical stars indicate statistical significance for the same strain under different conditions. Two-way ANOVA followed by Tukey’s multiple comparisons test was used.

We further investigated the role of the three AmpD paralogs and the two beta-lactamases in *C. violaceum* (Figure S3). The growth of all mutants was similar to that of the WT strain, except for the triple mutant (Δ*ampD1D2D3*), which showed slower growth (Figure S3A). In the viability assay, the Δ*ampD1*, Δ*ampD1D2* and Δ*ampD1D3* mutants showed 1 to 2 log reductions in CFU, if compared with the WT strain (Figure S3B), with *ampD1* having more dead cells (around 55%, Figure S3C). Curiously, the Δ*ampD1D2D3* triple mutant had no loss of viability, despite its slower growth (Figure S3A, B and C) but their colonies were smaller and displayed a faded purple color. The beta-lactamase mutants had no changes in fitness (Figure S3A, B and C). Taken together, these data reveal that AmpD1 appears to be the most important amidase for survival in the stationary phase (20 hours of culture) and that only the deletion of the three amidase genes impacts the growth of *C. violaceum*. Also, due to the importance of amidases in the correct functioning of the PGN, all the null mutants were evaluated for biofilm formation using the crystal violet staining method. All the strains harboring the *ampD1* deletion showed a reduction in biofilm formation (Figure S3D). The same was observed for violacein production, except for Δ*ampD1D3* (Figure S3E). The three beta-lactamase mutants had no changes in biofilm formation or violacein production (Figure S3D and E).

## DISCUSSION

In this work, we demonstrate that *C. violaceum* harbors two active and inducible chromosomally encoded beta-lactamases, AmpC and CphA, which confer resistance to distinct beta-lactam antibiotics. Moreover, we provide evidence that mutations in the amidase AmpD1, but not in its paralogs AmpD2 and AmpD3, consist of a major pathway toward stable overexpression of *ampC* and *cphA* and increased resistance to beta-lactams.

Our results using null *ampC* and *cphA* mutants in *C. violaceum* (Figure 1 and Table 1) and heterologous expression of these genes in *E. coli* (Figure S1) indicate that CphA is a metallo-beta-lactamase conferring resistance to carbapenems, while AmpC is a broad-spectrum beta-lactamase that confers resistance to penicillins and cephalosporins. These findings agree with the activity spectrum against beta-lactams described for AmpC and CphA in other bacteria (6–10). The complementary activities of these two beta-lactamases rise a concern because of their widespread co-occurrence in *Chromobacterium* species (27). Interestingly, a broad-spectrum KPC-like (class A) beta-lactamase was found in *C. piscinae*, *C. haemolyticum*, and *Chromobacterium sp.* C-61 (35, 36), but not in *C. violaceum*.

We investigated inducers and genetic mutations associated with beta-lactam resistance due to changes in beta-lactamase expression levels in *C. violaceum*. Our expression data indicate that *ampC* and *cphA* are induced in response to beta-lactams that vary between weak and strong inducers (Figure 2). These results are consistent with the expression pattern of chromosomally encoded beta-lactamases in other bacteria (12, 30, 31). We were able to isolate spontaneous mutants (SMs) in the presence of ceftazidime (CAZ), a clinically relevant third generation cephalosporin; 13 of the SMs were further characterized. These SMs showed high MIC values for CAZ (Table 2), increased resistance to most tested beta-lactams (Figure 3), hyperexpression of *ampC* and *cphA* (12 SMs) (Figure 4), and they carry mutations of different types in the *ampD1* gene (11 SMs) (Table 3; Figure 6). These data indicate that mutation in *ampD1* is an important route for the emergence of beta-lactam-resistant strains overexpressing AmpC and CphA beta-lactamases in *C. violaceum*. Indeed, mutations in AmpD amidases, which are enzymes involved in PGN recycling, have been associated with increased expression of beta-lactamases and resistance to beta-lactam antibiotics in clinical isolates of different bacteria, such as *P. aeruginosa*, *Burkholderia cenocepacia*, *Citrobacter freundii*, *Enterobacter cloacae*, and *E. coli* (11–13, 32, 33, 37–41).

Because we found three AmpD paralogs in *C. violaceum* (Figure 5), and mutations were detected in *ampD1* but not in *ampD2* or *ampD3* in the SM isolates (Table 3; Figure 6), we constructed null mutants that lack one, two or all three *ampD* genes to understand their individual contribution to beta-lactam resistance. We provide the following evidence that AmpD1 has a preponderant role over AmpD2 and AmpD3 in mediating the regulation and resistance to beta-lactams in *C. violaceum*: (i) null mutation in *ampD1*, but not in *ampD2* or *ampD3*, increased the resistance to beta-lactams (Figure 7; Table 4) and induced high expression of *ampC* and *cphA* beta-lactamases (Figure 9); (ii) except for SM59, *ampD1* rescued the susceptibility to CAZ when introduced into all the other 10 SM isolates that harbored a point mutation in *ampD1* (Figure 8). These findings differ from those that have been described in *P. aeruginosa,* in which sequential inactivation of its three amidases in a triple mutant was required to obtain high levels of *ampC* and resistance to beta-lactams (42–44). Remarkably, AmpD1 from *C. violaceum* has an acetyltransferase domain (ACTF_1; PF00583; EC 2.3.1) in its N-terminus belonging to the GNAT (N-acetyltransferases-like-Gcn5) family (Figure 5). This extra domain is not found in AmpDs characterized in other bacteria (Figure 6). Our preliminary *in silico* analyses suggest that amidases with this domain architecture are restricted to bacteria phylogenetically close to *C. violaceum* (genera *Aquitalea spp.*, *Chromobacterium spp.*, *Gulbenkiania spp.*, *Pseudogulbenkiania spp.*, and *Vogesella spp*). Further studies are needed to understand the role of the acetyltransferase domain found in the AmpD1 amidase of *C. violaceum*.

Mutations conferring antibiotic resistance can cause reduction in bacterial fitness (44, 45). Among the 13 SM isolates, only SM1 showed a delay in the growth curves, while the other isolates showed a minor loss of viability (Figure S2). With respect to the null *ampD* mutants, the mutant strains with *ampD1* deletion showed a moderate reduction in viability, biofilm, and violacein production. Only the triple mutant Δ*ampD1D2D3* showed a slower growth (Figure S3). These minor fitness losses in the absence of the three AmpD amidase paralogs suggest the presence of other amidases in *C. violaceum*. Indeed, a potential fourth amidase in *C. violaceum* (CV_3822) has an amidase-3 domain (PF01520) (Figure 5) that in *E. coli* is found in the amidases AmiA, AmiB and AmiC and which are involved in cell division (46, 47). More studies are required to understand the role of the *C. violaceum* amidases in cell division, growth, and survival.

In this work, we demonstrate that the beta-lactamases AmpC and CphA contribute to beta-lactam resistance in *C. violaceum*. We identified novel mutations in the unusual amidase AmpD1 that cause stable overexpression of *ampC* and *cphA*, providing new insights into the molecular mechanisms of beta-lactam resistance mediated by chromosomally encoded beta-lactamases. Altogether, our data offer an explanation for the limited effectiveness of many beta-lactams in treating *C. violaceum* infections and may pave the way for the development of more effective alternatives. Future studies should search for the transcription factors mediating the regulation of *ampC* and *cphA* and their relationship with the AmpD1 pathway described in this work. Moreover, genome sequencing of the SM1 and SM2 isolates, which were highly resistant to beta-lactams and did not harbor *ampD1* mutations, could reveal alternative mechanisms of beta-lactam resistance in *C. violaceum*.

## MATERIALS AND METHODS

### Bacterial strains, plasmids, and growth conditions

The bacterial strains and plasmids used in this work are indicated in Supplementary Table 1. *E. coli* and *C. violaceum* strains were cultured in Luria-Bertani (LB) or Mueller Hinton (MH) medium at 37 °C. When required, the media were supplemented with kanamycin (50 μg/mL), tetracycline (10 µg/mL) or ampicillin (100 μg/mL).

### Construction of *C. violaceum* mutant and complemented strains

In-frame null mutant strains were generated by allelic exchange mutagenesis, as previously described (22, 48). The flanking regions of the gene to be deleted were amplified by PCR using specific primers (Supplementary Table 2) and cloned into the suicide vector pNPTS138. The transconjugants were plated on LB 16% sucrose, and the null mutants were confirmed by PCR. For genetic complementation, the genes were amplified by PCR using specific primers (Supplementary Table 2), and cloned into the low copy number vector pMR20 (22, 48). All resulting constructs (for deletion or complementation) were transferred into the *C. violaceum* target strains by conjugation.

### Isolation of ceftazidime-resistant mutants

Spontaneous mutants resistant to ceftazidime were isolated as previously described (12). Briefly, an inoculum of *C. violaceum* ATCC 12472 WT strain from a single colony was incubated overnight in MH broth at 37 °C under agitation. A total of 100 μL of the inoculum were spread on MH agar plates containing increasing concentrations of ceftazidime (40, 80, 160, and 320 μg/mL) and the plates were incubated for 24 hours at 37 °C. Visible and isolated colonies obtained from the 80 μg/mL and 160 μg/mL MH plates were picked twice in antibiotic-free MH plates, and stored in 20% glycerol at – 80 °C.

### Antibiotic susceptibility tests

The MIC values were measured by using the agar dilution method according with the recommendations of the Clinical & Laboratory Standards Institute (28, 49). Briefly, fresh colonies grown on Mueller Hinton (MH) agar were suspended to OD_600 nm_ 0.1, and diluted to OD_600 nm_ 0.01. Drops of 2 μL of this bacterial inoculum (∼10^4^ CFU) were plated on MH agar prepared with or without different serial concentrations of the antibiotics. Bacterial growth was evaluated after incubation at 37 °C for 20 hours.

The resistance profile for several beta-lactam antibiotics was evaluated by disk diffusion assays performed as described by CLSI (28, 50). Briefly, fresh colonies grown on MH agar were suspended in 1X PBS to OD_600 nm_ 0.1. The bacterial suspension was seeded using a sterile swab on MH plates and disks with beta-lactam antibiotics (Table S3) (BD BBL Sensi-Disc or Thermo Scientific Oxoid) were applied on the top. After incubation at 37 °C for 20 hours, the inhibition halos were measured. In the MIC and disk diffusion assays, the *E. coli* ATCC 25922 strain was used as a control.

To detect metallo-beta-lactamase (MBL) activity in *C. violaceum*, we performed the modified carbapenem inactivation method (mCIM) and the EDTA-carbapenem inactivation method (eCIM) (28). Briefly, the bacterial strains were grown in trypticase soy broth (TSB) without (mCIM) or with 5 mM EDTA (eCIM). A disk with 10 μg imipenem (IPM) was added, and the cultures were incubated under agitation at 37 °C for 4 hours. The IPM disks were removed from the cultures and applied on the top of a *E. coli* ATCC 25922 indicator strain seeded on MH agar. The plates were incubated for 18-24 hours at 37 °C. The *K. pneumoniae* ATCC BAA 1705 strain was used as a control.

### DNA sequencing

The entire *ampD* genes (CV_0566/*ampD1* - 1259 pb; CV_1309/*ampD2* - 1225 pb; and CV_3031/*ampD3* - 1195 pb) were amplified by colony PCR from *C. violaceum* WT and SM isolates and sequenced in both strands. The purified PCR products were quantified in a NanoDrop spectrophotometer (Thermo Scientific). Sanger DNA sequencing reactions were prepared using the BigDye Terminator V3.1 kit (Applied Biosystems), PCR products (40 ng), and suitable primers (Supplementary Table 2), according to the manufacturer protocol. The reactions were precipitated by adding 75% isopropanol and washed two times with 70% ethanol. DNA sequencing was carried out on an ABI 3500XL (Applied Biosystems).

### Transcriptional *lacZ* fusions and beta-galactosidase assays

The upstream regions of the *ampC* and *cphA* genes were amplified by PCR (Supplementary Table 2) and cloned into the pGEM-T easy plasmid (Promega). The inserts were subcloned into the pRK*lacZ*290 vector to obtain transcriptional fusions of the *lacZ* gene. *C. violaceum* strains containing the reporter plasmids were grown to OD_600 nm_ 1 in LB. The cultures were divided and untreated or treated with following antibiotics: ceftazidime (CAZ) 30 μg/mL, cefoxitin (FOX) 10 μg/mL, ampicillin (AMP) 100 μg/mL, imipenem (IPM) 1 μg/mL, kanamycin (KAN) 50 μg/mL, gentamicin (GEN) 40 μg/mL, streptomycin (STR) 20 μg/mL, nalidixic acid (NAL) 20 μg/mL, ciprofloxacin (CIP) 1 μg/mL, tetracycline (TET) 3 μg/mL, polymyxin B (PMB) 10 μg/mL μg/mL, and rifampin (RIF) 10 μg/mL. After 30 minutes, aliquots of the cultures (100 μL) were assayed for beta-galactosidase activity using a previously described protocol (48, 51).

### RNA extraction and expression analysis by RT-qPCR

*C. violaceum* strains were cultured in LB at 37 °C under agitation until OD_600 nm_ 1. The cultures were divided and untreated or treated with 100 μg/mL ampicillin (AMP) for 30 minutes. Total RNA was extracted in TRIzol reagent (Invitrogen) and purified using the Direct-zol RNA miniprep plus kit (Zymo), according to the manufacturer’s instructions. The quantity and quality of the RNA was assessed on a denaturing agarose gel and on a Nanodrop spectrophotometer (Thermo Scientific). The cDNA was synthesized using the High-Capacity cDNA Reverse Transcription kit (Applied Biosystems). The qPCR (quantitative PCR) reactions were performed with the PowerUp™ SYBR™ Green Master Mix kit (Applied Biosystems), 10 ng of cDNA and 0.5 μM of specific primers (Supplementary Table 2), using the QuantStudio 3 thermal cycler (Applied Biosystems). The results were analyzed using the QuantStudio Design & Analysis v1.5.2 software with the 2^−ΔΔC*t*^ method (52). Data were normalized to the endogenous *minD* (CV_3376) gene and a reference condition (wild-type strain *C. violaceum* in LB).

### Growth curves

Bacterial growth in LB was monitored over time using absorbance measurements at 600 nm on the BioTek Epoch 2 (Agilent). An overnight pre-inoculum in LB was adjusted to OD_600 nm_ 0.01 in LB, and 150 μL of this dilution were added to a 96-well flat-bottomed microtiter plate. Plates were incubated under 425 CPM orbital shaking (3 mm diameter) at 37 °C for 18 hours. Measurements were taken every hour. These assays were carried out in biological triplicate.

### Survival assays

To calculate colony-forming units per milliliter (CFU/mL), *C. violaceum* cultures were diluted to OD_600 nm_ 0.01 in 5 mL of LB, and grown in a shaker at 37 °C, 250 RPM for 20 hours. After centrifugation, the cultures were serially diluted (1:10) in 1X phosphate buffered saline (PBS) and plated on LB agar. The colonies were counted after incubating for 20 hours at 37 °C. Alternatively, bacterial survival was evaluated using the LIVE/DEAD BacLight Bacterial Viability kit (Thermo Fisher Scientific), according to the manufacturer’s instructions. Cultures were grown for 20 hours in LB, washed three times with 0.85% NaCl (w/v), and the cells were stained with 2X LIVE/DEAD solution in a 1:1 ratio for 15 minutes. Samples were prepared on an agarose pad and visualized using a Leica TCS SP5 confocal microscope. Images of randomly obtained fields were analyzed using ImageJ software.

### Static biofilm assay

Biofilm formation was quantified using the crystal violet staining method (48). The *C. violaceum* strains were grown from an OD_600 nm_ 0.01 in LB in glass tubes under incubation at 37 °C for 24 hours without shaking. The culture was discarded, and 0.1% crystal violet was added for 15 minutes. The tubes were washed, dried, and 33% acetic acid was added. After 1 hour at room temperature, the biofilm was quantified by OD_600 nm_, and normalized by the growth of the culture.

### Analysis of violacein production

Overnight pre-inoculum of *C. violaceum* cultures in LB was diluted at OD_600 nm_ 0.01 in 2 mL of LB, and incubated at 37 °C for 24 hours under 250 RPM agitation. After that, 500 μL of each culture were mixed with 500 μL of 100% acetone, and vortexed for 30 seconds. Violacein was quantified from the organic phase by measuring absorbance at 575 nm, and the values were normalized by the growth of the culture.

## ACKNOWLEDGMENTS

This study was financed, in part, by the São Paulo Research Foundation (FAPESP; process numbers 2021/06894-0 and 2021/10577-0), Brasil, and Fundação de Apoio ao Ensino, Pesquisa e Assistência do Hospital das Clínicas da FMRP-USP (FAEPA). I.H. lab has support from Fundação para a Ciência e a Tecnologia (FCT) through the Research Units CFE (https://doi.org/10.54499/UIDB/04004/2020) and Associate Laboratory TERRA (https://doi.org/10.54499/LA/P/0092/2020). During this work, L.G.L. was supported by fellowships from FAPESP (2021/01911-3 and 2022/07135-8) and CAPES (Coordenação de Aperfeiçoamento de Pessoal de Nível Superior). C.E.M.N. was supported by FAPESP fellowship (2018/02465-4). J.F.S.N. is Research Fellow from CNPq (Conselho Nacional de Desenvolvimento Científico). We would like to thank Marcia Triunfol at Publicase International for her assistance in editing this manuscript.

